# Intracranial substrates of meditation-induced neuromodulation in amygdala and hippocampus

**DOI:** 10.1101/2024.05.10.593445

**Authors:** Christina Maher, Lea Tortolero, Daniel D. Cummins, Adam Saad, James Young, Lizbeth Nunez Martinez, Zachary Schulman, Lara Marcuse, Allison Waters, Helen S. Mayberg, Richard J. Davidson, Ignacio Saez, Fedor Panov

**Affiliations:** Nash Family Department of Neuroscience and the Friedman Brain Institute, Icahn School of Medicine at Mount Sinai, New York, NY, USA; Nash Family Center for Advanced Circuit Therapeutics, Icahn School of Medicine at Mount Sinai, New York, NY, USA; Department of Psychiatry, Icahn School of Medicine at Mount Sinai, New York, NY, USA; Department of Neurosurgery, Icahn School of Medicine at Mount Sinai, New York, NY, USA; Department of Neurology, Icahn School of Medicine at Mount Sinai, New York, NY, USA; Departments of Psychology and Psychiatry, University of Wisconsin–Madison, Madison, WI, USA

**Keywords:** loving-kindness meditation, amygdala, hippocampus, intracranial electrophysiology

## Abstract

Meditation is an accessible mental practice associated with emotional regulation and well-being. Loving-kindness meditation (LKM), a specific sub-type of meditative practice, involves focusing one’s attention on thoughts of well-being for oneself and others. Meditation has been proven to be beneficial in a variety of settings, including therapeutical applications, but the neural activity underlying meditative practices and their positive effects are not well understood. In particular, it’s been difficult to understand the contribution of deep limbic structures given the difficulty of studying neural activity directly in the human brain. Here, we leverage a unique patient population, epilepsy patients chronically implanted with responsive neurostimulation device that allow chronic, invasive electrophysiology recording to investigate the physiological correlates of loving-kindness meditation in the amygdala and hippocampus of novice meditators. We find that LKM-associated changes in physiological activity specific to periodic, but not aperiodic, features of neural activity. LKM was associated with an increase in γ (30-55 Hz) power and an alternation in the duration of β (13-30 Hz) and γ oscillatory bursts in both the amygdala and hippocampus, two regions associated with mood disorders. These findings reveals the nature of LKM-induced modulation of limbic activity in first-time meditators.

**SIGNIFICANCE STATEMENT:** We leverage rare chronic, invasive electrophysiology recordings while participants engage in loving-kindness meditation to demonstrate that meditation induces neural changes in beta and γ activity in the amygdala and hippocampus of novice meditators. These results build on previous findings in experienced meditators and reveal meditation’s potential for noninvasive neuromodulation of neural mechanisms associated with emotional regulation and mood disorders.

## INTRODUCTION

Meditation is a set of mental techniques aimed at cultivating well-being, which require honing attentional skills related to emotional regulation (Chambers et al., 2009; Wadlinger & Isaacowitz, 2006). Numerous studies have proven that meditation can improve mental well-being in both a population-based setting (Shapiro et al., 1998), and potentially improve psychiatric diseases such as anxiety and depression (Wielgosz et al., 2019). In concert with its clinical effects, meditation has been shown to change both brain electrophysiology and functional neuroimaging (Boccia et al., 2015; Kaur & Singh, 2015). Recent literature has parceled out primary categories of meditation, thus allowing rigorous scientific evaluation of specific practices (Dahl et al., 2015). Loving-kindness meditation (LKM) is a technique within the constructive meditation family, in which practitioners actively focus their attention on cultivating positive thoughts of well-being for others. Preliminary work has suggested varying forms of meditation may share common effects on brain electrophysiology (Braboszcz et al., 2017): this remains in important area for potential study. Loving-kindness meditation may have therapeutic potential through the cultivation of positive emotion (Petrovic et al., 2024), but its underlying neural correlates are not well known, especially in deep brain areas involved in emotional regulation.

Functional and structural magnetic resonance imaging (MRI) studies have demonstrated changes in both the amygdala and the hippocampus from continued LKM practice (Desbordes et al., 2012; Leung et al., 2013). EEG studies, in contrast, have shown increased γ activity during meditation (Braboszcz et al., 2017; Leung et al., 2013), including during LKM in experienced meditators (Lutz et al., 2004). BOLD-fMRI signal has been shown to correlate with γ activity (Lachaux et al., 2007), suggesting that these processes could be related, but the nature of human amygdala and hippocampal activity during LKM, including a physiologically detailed and anatomically precise depiction of neural changes during meditative practice, remains to be directly ascertained, and several questions remain. For example, whether LKM is specifically associated with oscillatory processes rather than an overall excitation/inhibition profile modulation in associated neural circuits has not been determined. In addition, the involvement of other frequency bands (e.g. β = 12-30Hz) in limbic regions remains unclear, despite their role in emotional regulation and mood processes (Alagapan et al., 2023; Sani et al., 2018; Young et al., 2024). Meditation has been associated with modulating local neuronal activity (Lutz, Brefczynski-Lewis, et al., 2008; Lutz et al., 2004; Lutz, Slagter, et al., 2008), measured by oscillatory power estimates (Donoghue et al., 2020), and decreased population synchrony within and between discrete regions (Irrmischer, Houtman, et al., 2018; Irrmischer, Poil, et al., 2018), measured by the duration of rhythmic, oscillatory episodes (Kosciessa et al., 2020).

Therefore, despite its importance, the anatomically precise and physiologically detailed neural basis of meditative practice remains to be determined, especially in deep brain areas that are inaccessible to non-invasive electrophysiological recording methods. This, in turn, limits our understanding of the neural changes associated with the positive impacts of meditative practices and the development of generalizable insights that may be useful for therapeutical development. Neurosurgical interventions for the management of epilepsy allow recording from such areas in humans, including recording local field potentials (LFPs) capturing circuit activity directly via intracranial depth leads with great electrophysiological detail and signal to noise ratio (von Ellenrieder et al., 2012), and providing a unique opportunity to study the neural basis of human behavior and thought. However, the most common setting for these invasive recordings, during drug resistant epilepsy (DRE) patients’ hospitalization (e.g. in the Epilepsy Monitoring Unit) are not ideal for the study of meditation because of their perioperative nature and the lack of an adequate environment for calm meditative practices. In contrast, responsive neurostimulation (RNS) system (NeuroPace Inc.) allows chronic electrophysiologic brain recordings from implanted regions, frequently from the mesial temporal lobe structures of the hippocampus and amygdala (Arcot Desai et al., 2019), during the patients’ daily life after surgery. This allows combining intracranial recordings with the practice of meditation in a controlled setting providing adequate environmental conditions for the practice of meditation.

Patients implanted with the RNS device can move around freely while continuous iEEG activity is recorded as local field potentials (LFPs). Recordings made with the RNS also offer high-quality data from deep brain structures, implicated in emotional regulation, such as the mesial temporal lobe (Qasim et al., 2023). This is a major advantage in contrast to meditation studies using scalp surface EEG, which have significantly lower signal-to-noise ratio and do not allow the high simultaneous spatial and temporal resolution of RNS iEEG recordings (Parvizi & Kastner, 2018). Patients implanted with the RNS are thus ideal candidates for investigating the neural correlates of naturalistic behavior, such as meditation (M. Aghajan et al., 2017; Maoz et al., 2023). In addition, DRE patients often suffer from psychiatric comorbidities including depression and anxiety (Josephson et al., 2017), which provides an opportunity to study the relationship between intracranial activity and comorbid state, as well as the potential modulation during meditation.

We therefore explored changes in neural oscillatory activity associated with LKM within the amygdala and hippocampus using iEEG in DRE patients chronically implemented with an RNS device. Our findings show that first-time LKM modulates frequency-specific power (Fig 5) and duration (Fig 6) of oscillatory events in the hippocampus and amygdala. The selective nature of these findings in regard to the modulation of periodic, but not aperiodic, features of the neural signal reveals potential biomarkers by which LKM, a readily accessible therapeutic technique, noninvasively modulates physiological processes associated with mood regulation (Alagapan et al., 2023; Sani et al., 2018; Young et al., 2024) even in first-time meditators.

## RESULTS

Participants included eight neurosurgical patients with drug-resistant epilepsy who were chronically implanted with the NeuroPace Responsive Neurostimulation System (RNS). Participants completed the present study in Mount Sinai’s Quantitative Biometric Laboratory (Q-Lab), designed to provide patients with a relaxing environment to receive therapeutic treatment free from typical distractors associated with a hospital setting and therefore highly conducive to engaging in meditative practice (Fig. 1B). Participants were self-reported novice meditators prior to the present study and completed a 5-minute audio-guided instruction (baseline) followed by 10 minutes of audio-guided LKM (See Methods). We analyzed LFPs by creating bipolar derivations between the two most anterior contacts (typically located in the amygdala and anterior hippocampus) and the two most posterior contacts (the middle-posterior hippocampus). Therefore, we collected two bipolar channels (one anterior pair and one posterior pair) per hemisphere implanted with RNS. Six patients had bilateral RNS implantations, and two patients had unilateral RNS implantation (left hemisphere). All participants included in the present study had at least one contact in either the amygdala or hippocampus (count: amygdala = 14 electrodes/13 bipolar channels; hippocampus = 36 electrodes/14 bipolar channels; Fig 2A). Anatomical localization of electrodes was determined by co-registering high-resolution post-operative CT scans with pre-operative magnetic resonance images (MRI) (See Methods). Although RNS implantation occurs in DRE patient’s presumptive seizure onset zone, the leads which are composed of 4 contacts typically span just over 30 mm of tissue (1.5mm in length for each contact and 10mm between contact centroids). This results in a proportion of data recorded from likely normative tissue (M. Aghajan et al., 2017; Maoz et al., 2023). To mitigate the influence of interictal noise in the data, we implemented a data preprocessing approach mentioned in previous publications (Saez et al., 2018). Briefly, following visual inspection of all channel data, we confirmed the absence of any stimulation artifact and eliminated any data meeting clinical criterion for interictal discharge (Marcuse, Lara et al., n.d.). This resulted in discarding approximately 6% of the data. The proportion of data across amygdala and hippocampus bipolar channels (n=27) in which interictal or noisy activity was detected was not significantly different between conditions (baseline and meditation; p > 0.05 for all channels, Pearson’s chi-square goodness-of-fit test).

**Figure 1.**
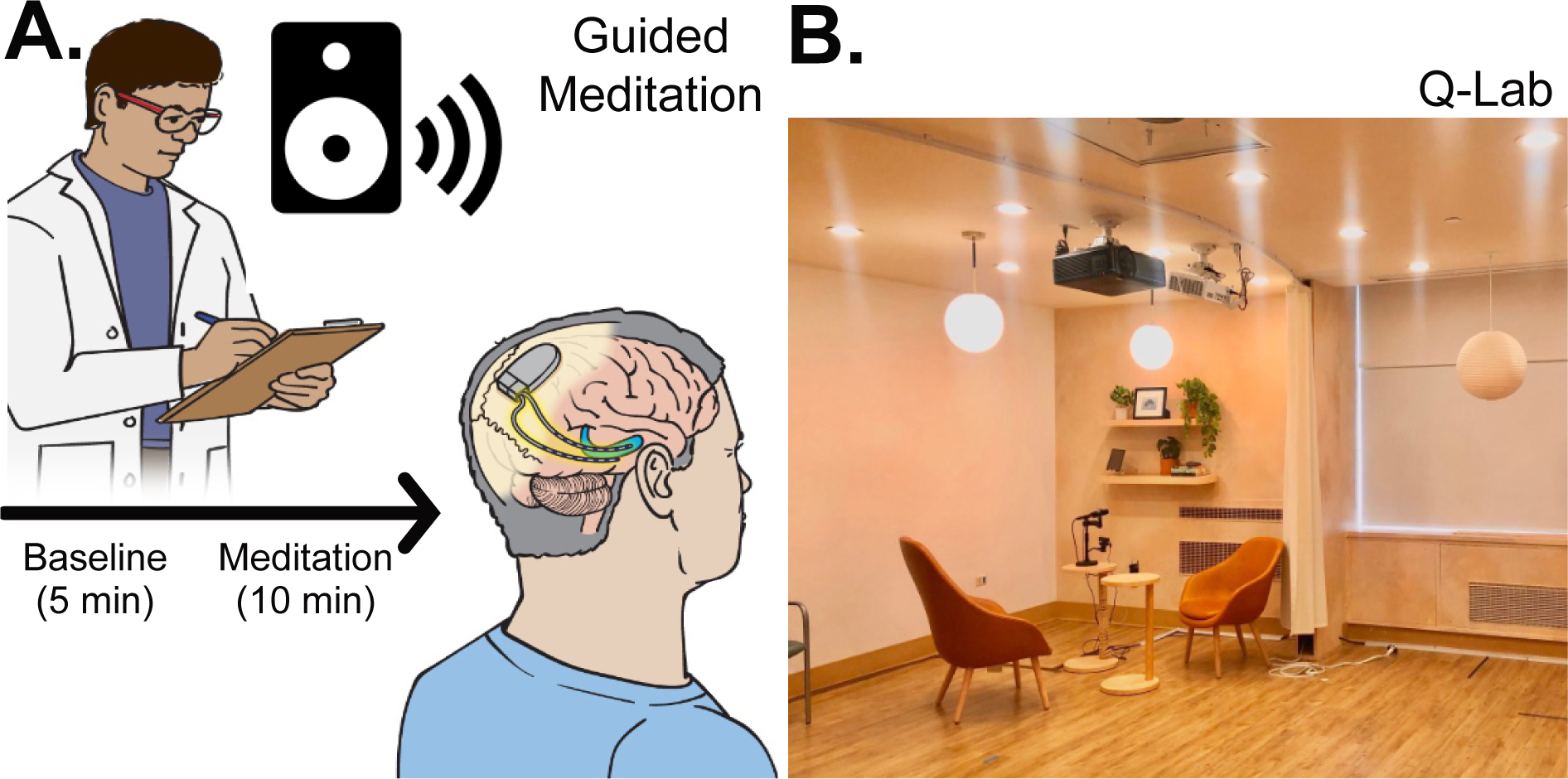
Behavioral methods. A. Experimental design. Subjects (n=8) completed a LKM paradigm consisting of 5 minutes of audio-guided instruction (baseline) and 10 minutes of audio-guided LKM. **B. Experimental setting.** The experimental paradigm was administered in Mount Sinai West’s Quantitative Biometrics Laboratory (Q-Lab), a dedicated, immersive research environment designed to provide participants with a restorative space to participate in this experiment.

**Figure 2.**
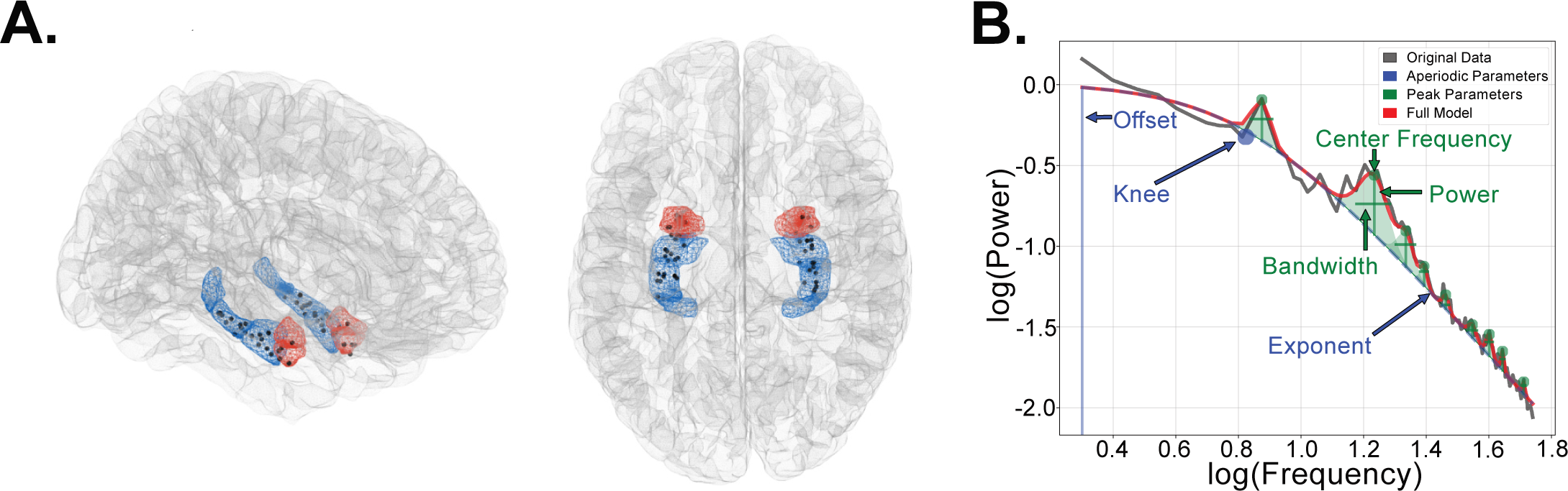
Neural methods. A. Anatomical reconstruction showing the hippocampal and amygdala location of RNS contacts. Depicted is the placement of 50 NeuroPace RNS electrodes in amygdala (blue) and hippocampus (red) across 8 participants. Each black dot corresponds to one electrode (amygdala = 14 electrodes/13 bipolar channels; hippocampus = 36 electrodes/14 bipolar channels). **B. Fitting Oscillations and One Over F approach (FOOOF).** We characterized local field potential (LFP) activity from bipolar channels recorded from depth electrodes in the amygdala and hippocampus. We used the Fitting Oscillations and One Over F approach (FOOOF) to separately characterize aperiodic (i.e. 1/f background activity) from oscillatory neural activity. Depicted is an example power spectrum from one hippocampal channel (gray trace) overlaid by the FOOOF fitted model (red trace) parameterized to extract aperiodic (blue trace) and periodic (green trace) spectral features between 2 and 55 Hz. The aperiodic components are characterized by the offset, knee and exponent. Periodic components are assigned to a canonical frequency band (δ=2-4Hz, θ=4-8Hz, α=8-13Hz, β=13-30Hz, γ=30-55Hz) depending on their center frequency, and their power and bandwidth estimated.

### LKM was not accompanied by changes in aperiodic neural activity

We set out to identify whether meditation was accompanied by changes in neural activity by comparing LFP activity patterns between active control (learning about meditation) and LKM epochs. Given the largely temporally unresolved nature of this data, we chose to focus our analysis on the power profile throughout a single epoch during the meditation block, examining both aperiodic (1/f profile) and periodic (i.e. oscillatory) neural activity on each bipolar channel. We parameterized the aperiodic and periodic features of the power spectra using the Fitting oscillations and one over-f method (FOOOF; see Methods and Fig. 2) (Donoghue et al., 2020). The FOOOF model fit was performed for each channel’s data in each condition (baseline and meditation). The aperiodic components include a knee parameter, which accounts for an often-observed bend in the 1/f profile, and offset and exponent, which reflect the y-intersect and rate of decay of the 1/f profile, respectively. Together, these aperiodic components capture broadband shift in the 1/f profile often ascribed to changes in excitatory/inhibitory balance, different and separate from oscillations in individual frequency bands. In addition, oscillatory components were estimated for pre-defined frequency bands (δ=2-4Hz, θ=4-8Hz, α=8-13Hz, β=13-30Hz, γ=30-55Hz; see Methods).

We first sought out to identify whether acute LKM induced changes in the aperiodic features of the power spectra by comparing knee frequency, offset, and exponent separately between baseline and LKM epochs (Fig 4). We did not find significant differences between conditions in knee frequency, offset or exponent in either the amygdala or hippocampus (all p>0.05, two-sided paired-samples Wilcoxon signed rank test; Fig. 3), suggesting that meditation is not accompanied by general changes in the excitatory/inhibitory profile of either amygdala or hippocampus. Further, we found no significant differences in aperiodic components between amygdala and hippocampus (all p>0.05, two-sided Wilcoxon signed rank test; Supplementary Fig. 1).

**Figure 3.**
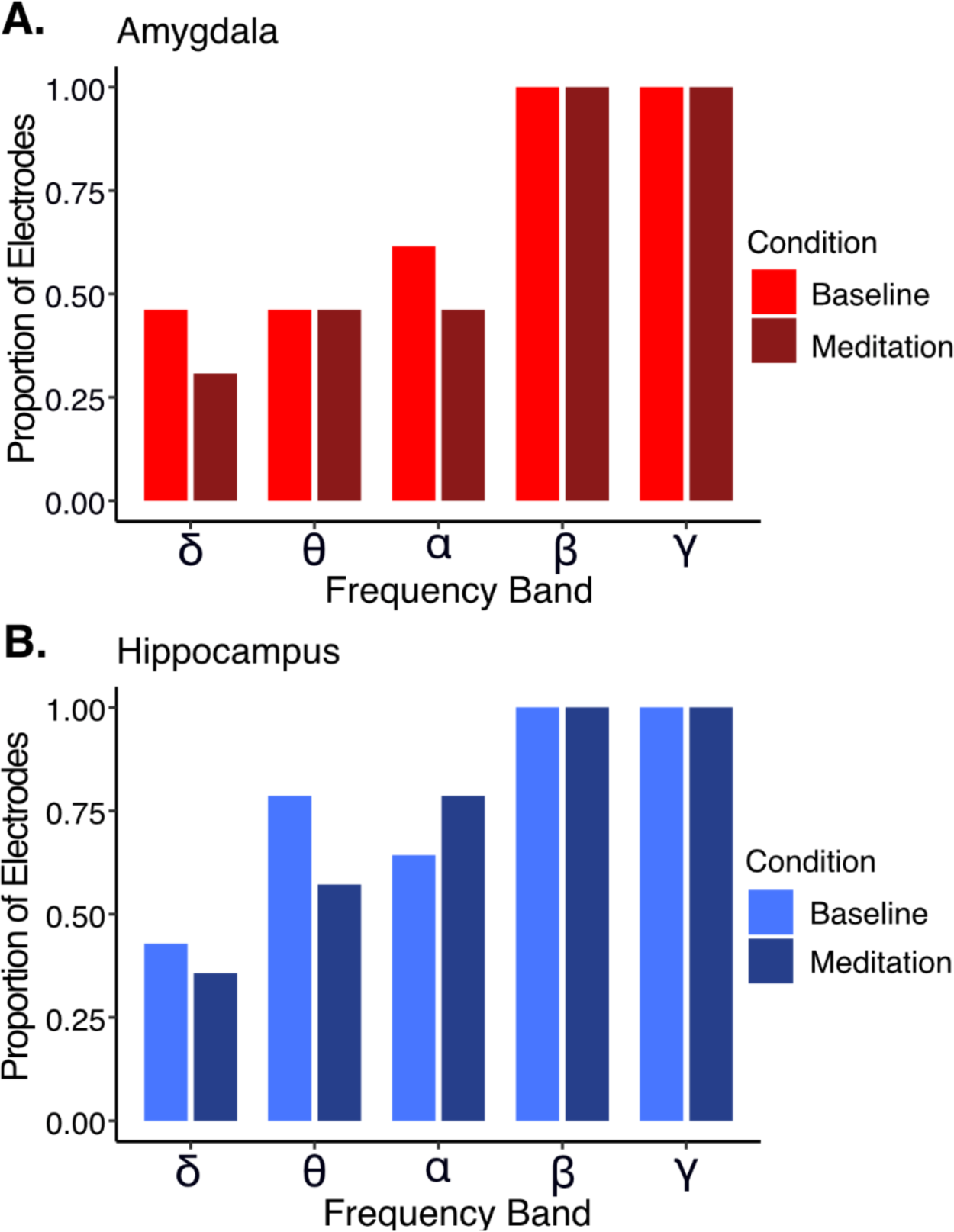
Proportion of channels showing neural oscillations varied across frequency bands. Plotted is the proportion of channels in which an oscillation was detected during baseline and meditation epochs in each frequency band (δ=2-4Hz, θ=4-8Hz, α=8-13Hz, β=13-30Hz, γ=30-55Hz) according to FOOOF power spectrum model fit. **A. Proportion of amygdala channels showing significant oscillatory activity across frequencies.** The proportion of channels showing significant modulation varied between ∼30% in δ (2-4Hz) and 100% in γ (30-55Hz), with greater proportions for higher frequencies. There were no differences in the proportion of amygdala channels (n=13 bipolar channels) in which oscillations were detected between baseline (light blue) and meditation (dark blue) epochs (all p>0.05, Fisher’s exact test). **B. Proportion of hippocampal channels showing significant oscillatory activity across frequencies.** The pattern of activation was like that observed in the amygdala, with the proportion of channels ranging between ∼30% in δ (2-4Hz) and 100% in γ (30-55Hz), with greater proportions for higher frequencies. There were no differences in the proportion of hippocampal channels (n=14 bipolar channels) in which oscillations were detected between baseline (light blue) and meditation (dark blue) epochs (all p>0.05, Fisher’s exact test).

**Figure 4.**
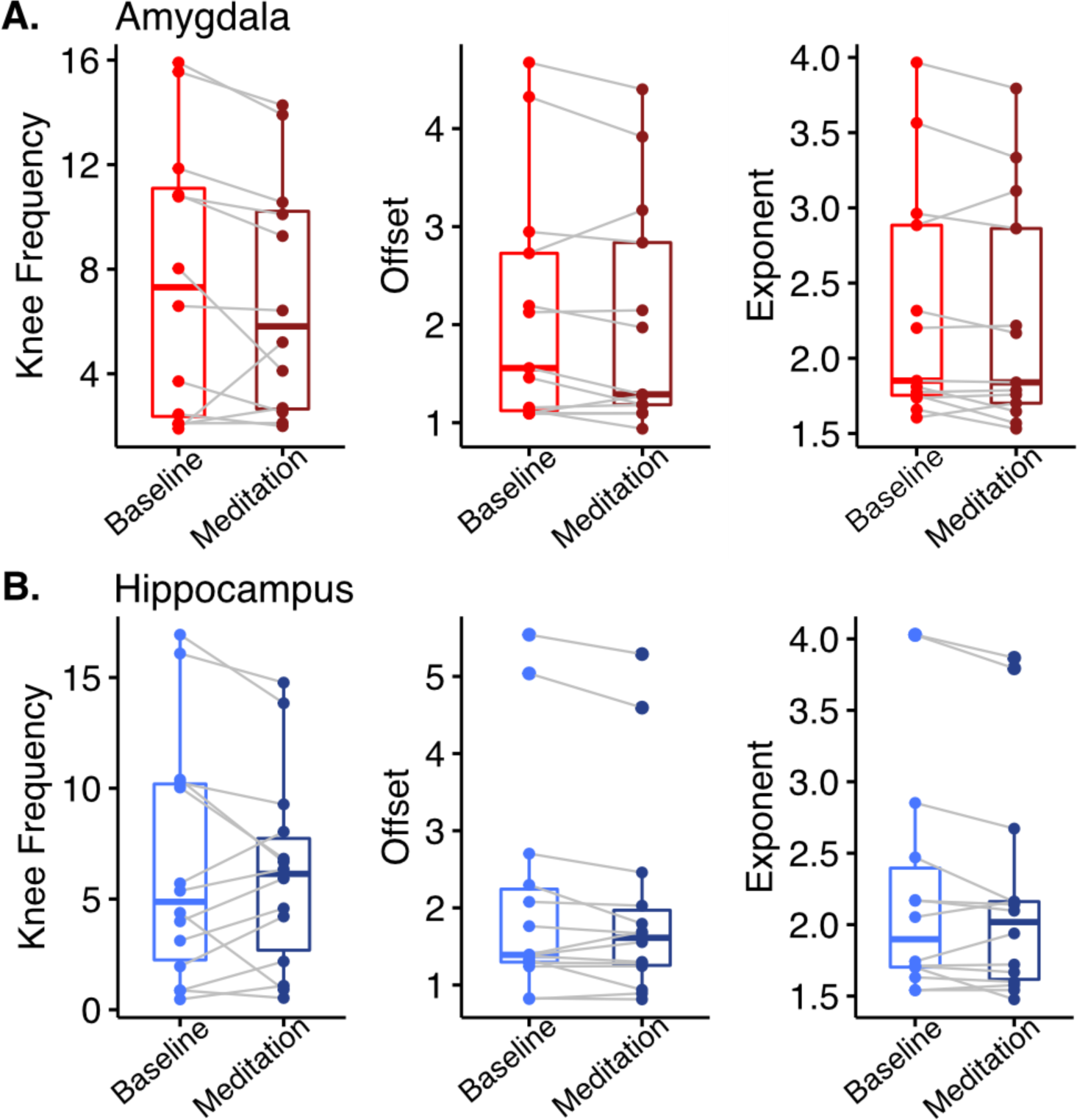
No difference in aperiodic neural components between baseline and meditation epochs. The aperiodic component of FOOOF model fit (knee frequency, offset, and exponent) were extracted and compared between conditions (FA meditation and baseline). **A. No difference in FOOOF aperiodic parameters in amygdala channels.** No significant differences in knee frequency, offset, or exponent of FOOOF model fit for amygdala channels (n=13 bipolar channels) between conditions (all p>0.05, Two-sided Paired sample Wilcoxon signed-rank test). **B. No difference in FOOOF aperiodic parameters in hippocampus channels.** No significant differences in knee frequency, offset, or exponent of FOOOF model fit for hippocampus channels (n=14 bipolar channels) between conditions (all p>0.05, Two-sided Paired sample Wilcoxon signed-rank test).

### LKM was not accompanied by changes in the proportion of electrodes showing oscillatory activity

Next, we evaluated whether LKM was accompanied by changes in oscillatory neural activity by examining power across frequencies in baseline and meditation conditions. We considered a significant oscillation to be present if at least one peak within pre-defined frequency bins (δ=2-4Hz, θ=4-8Hz, α=8-13Hz, β=13-30Hz, γ=30-55Hz) was detected by the FOOOF model fit. If no peak was found within a given frequency range for a channel, we considered the channel to not contain an oscillation in that band. Therefore, we started by quantifying the proportion of channels containing active oscillations across conditions. We observed differences in the proportion of active channels across frequencies between conditions in both amygdala and hippocampus, with higher frequencies containing a larger proportion of active channels than lower frequencies (Fig 4). The lowest proportion of active electrodes was in δ in both amygdala (30.8%) and hippocampus (35.7%); the highest was in β and γ in both regions (100%). However, we did not find significant differences in the proportion of active electrodes between conditions in any frequency bands (all p>0.05, Fisher’s exact test). Therefore, LKM was not associated with an increase in the proportion of channels showing significant oscillations. These data indicate that our ability to detect significant oscillations was strongly frequency-dependent, and that there was significant β and γ oscillatory activity in both baseline and in meditation conditions in the amygdala and hippocampus.

### LKM was associated with an increase in γ power

To further investigate whether LKM modulated oscillatory activity, we compared the amplitude, or power, of detected oscillations between conditions in both regions (Fig 5). To investigate between-condition differences in oscillatory power, we only considered channels in which at least one peak was detected in each frequency band (i.e. containing significant oscillatory activity). If more than one peak was found within a given frequency range, the average power of all detected peaks within the frequency band was computed to determine an average power score. We observed a significant increase in γ power during LKM in both the amygdala (p < 0.01, one-sided paired-samples Wilcoxon signed rank test, Fig 5A). and hippocampus (p < 0.001, one-sided paired-samples Wilcoxon signed rank test, Fig 5B). We did not observe concomitant changes in any other frequency bands (all p > 0.05, one-sided paired-samples Wilcoxon signed rank test, Fig. 5), indicating that this modulation was specific to the γ frequency band. Overall, these results reveal that LKM is accompanied by changes in oscillatory power in active channels, but not an increase in the proportion of active channels. Furthermore, these changes are specific to high-frequency activity (γ) and not present in lower frequencies. We did not find differences in baseline oscillatory power or the difference in oscillatory power from baseline to meditation between the amygdala and hippocampus (all p>0.05, two-sided Wilcoxon signed rank test; Supplementary Fig. 2a/3a).

### LKM was accompanied by an alteration in duration of β and γ bursts

The FOOOF method allowed us to investigate aperiodic and periodic components of brain activity but does not offer a way to quantify quick bursts of neural activity which may be important to facilitate switches in cognitive states (e.g. from baseline to LKM). To investigate this possibility, we applied the extended Better Oscillation detection (eBOSC) method (Kosciessa et al., 2020; Whitten et al., 2011), to investigate how meditation modulated the duration of oscillations across frequency bands (see Methods). eBOSC allows us to detect temporal windows with significant frequency-specific oscillations that surpass power and duration thresholds while accounting for background 1/f profile of the neural signal (Kosciessa et al., 2020). Further, recent work has implemented this method with hippocampal iEEG data collected using the RNS device, allowing us to compare our findings with previously published iEEG work (M. Aghajan et al., 2017). Using eBOSC, we calculated the proportion of time within a given experimental condition (baseline or LKM) in which a particular oscillation was detected. We averaged these proportions for each frequency in each of our pre-defined frequency bands and compared between conditions to identify whether LKM induced changes in the duration of oscillatory events (Fig. 6). We found that the meditative state was associated with a significant decrease in the duration of β oscillations in both the amygdala (p < 0.05, one-sided paired-samples Wilcoxon signed rank test, Fig. 6A). and hippocampus (p < 0.05, one-sided paired-samples Wilcoxon signed rank test, Fig. 6B). Further, we found a significant increase in the duration of γ oscillations during LKM in the amygdala (p < 0.01, one-sided paired-samples Wilcoxon signed rank test, Fig. 6A). Additionally, the amygdala displayed a significantly greater increase in the duration of γ oscillations during meditation compared to the hippocampus (p < 0.05, one-sided Wilcoxon signed rank test, Supplementary Fig 3b; all other frequency bands p>0.05). Overall, these results demonstrate a decrease in the amount of β events during meditation in both amygdala and hippocampus, and an increase in γ events in amygdala, suggesting an involvement of fast oscillatory events in meditative states. We did not find differences in baseline oscillatory duration between the amygdala and hippocampus (all p>0.05, two-sided Wilcoxon signed rank test; Supplementary Fig. 2b/3b).

## DISCUSSION

This study leveraged unique access to chronic ambulatory iEEG recordings in the amygdala and hippocampus during LKM. We directly examined the nature of neurophysiological activity during LKM in the amygdala and hippocampus, finding primary results of increased γ oscillations and decreased β duration during LKM. Further, changes in neural dynamics with meditation occurred in oscillatory activity, rather than in aperiodic neural mechanisms. Research on the effects of meditation on neural dynamics to date has been limited to fMRI and scalp EEG, due to rare access to iEEG in real world settings. While such approaches have advanced the field of meditation, iEEG research can augment such data with increased spatial and temporal resolution.

Meditative practices have existed for millennia in different traditions (Wynne, 2007) and allow humans to cultivate fundamental states of focus and emotional regulation (Manna et al., 2010). Meditation, perhaps independent of practice type, induces scalp EEG increases in γ activity (Braboszcz et al., 2017). In addition, meditation induces changes in BOLD-fMRI signal in the hippocampus and amygdala (Desbordes et al., 2012; Engström et al., 2010), which could correlate with changes in γ oscillatory activity (Lachaux et al., 2007). Our study directly addresses the nature of intracranial electrophysiology within the hippocampus and amygdala during meditation. Our focus on LKM comes from previous research that connects this emotionally laden contemplative practice to the amygdala (Desbordes et al., 2012). LKM, utilized in this study, stresses finding joy and sharing it with others (Dahl et al., 2015). Such constructive contemplative practices have been shown to induce positive emotions, which may lead to an increased sense of purpose and meaning in life (Fredrickson et al., 2008). Such training has also been shown to chronically modulate the amygdala, hence of specific interest in our cohort where contacts were situated in the basolateral amygdala in all cases (Desbordes et al., 2012). Yet, no experiments to date have attempted to record such changes in novice meditators acutely and intracranially. These findings complement the earlier results on increased γ oscillations in LKM practices in long-term meditators by showing specific increases in these fast oscillations in novice meditators when directly examining amygdala and hippocampus with iEEG.

### Amygdala and hippocampus γ power increases during LKM

By leveraging the spatiotemporal resolution afforded by iEEG, our results build on prior research on γ power increases associated with long-term LKM practice (Braboszcz et al., 2017; Ferrarelli et al., 2013; Garrison et al., 2015; Lutz et al., 2004; Lutz, Slagter, et al., 2008), extending these findings to first-time meditators and providing new insights into the nature of neural changes associated with meditation in an anatomically precise way. We observed an increase in amygdala and hippocampus γ power (Fig 5), whose correlation to fMRI-BOLD signal is well-established (Lachaux et al., 2007), suggesting LKM induces heightened activation of local neuronal ensembles within the amygdala and hippocampus. This meditation-induced neural modulation may be related to the known role in processing of emotional information in amygdala and hippocampus (Roberts-Wolfe et al., 2012), and possibly with related mental processes such as memory and attention. During LKM, participants are encouraged to actively retrieve positive autobiographical memories during LKM which induces amygdala and hippocampus activation (van Schie et al., 2019). The hippocampus plays a critical role emotional regulation and attention (Poskanzer & Aly, 2023; Zhu et al., 2019) alongside its fundamental roles memory consolidation and retrieval (Carr et al., 2011; Wiltgen et al., 2010), suggesting hippocampal activation may be related to the memory aspects of the meditative task. The amygdala, in turn, can direct attention toward emotionally significant stimuli (Desbordes et al., 2012)(Carr et al., 2011; Jacobs et al., 2012), serving a critical function in the bottom-up processing of emotionally-relevant information (Klinge et al., 2010; Peck & Salzman, 2014; Pessoa & Adolphs, 2010). We further identified an increase in the duration of γ oscillatory events during LKM compared to baseline in the amygdala, but not hippocampus (Fig 6). This may reflect the specific physiological demands of LKM meditation, wherein the hippocampus exhibits more transient γ bursts associated with memory retrieval (Lundqvist et al., 2024) while the amygdala exhibits more sustained, rhythmic γ activity to support emotional processing (Bocchio et al., 2017).

**Figure 5.**
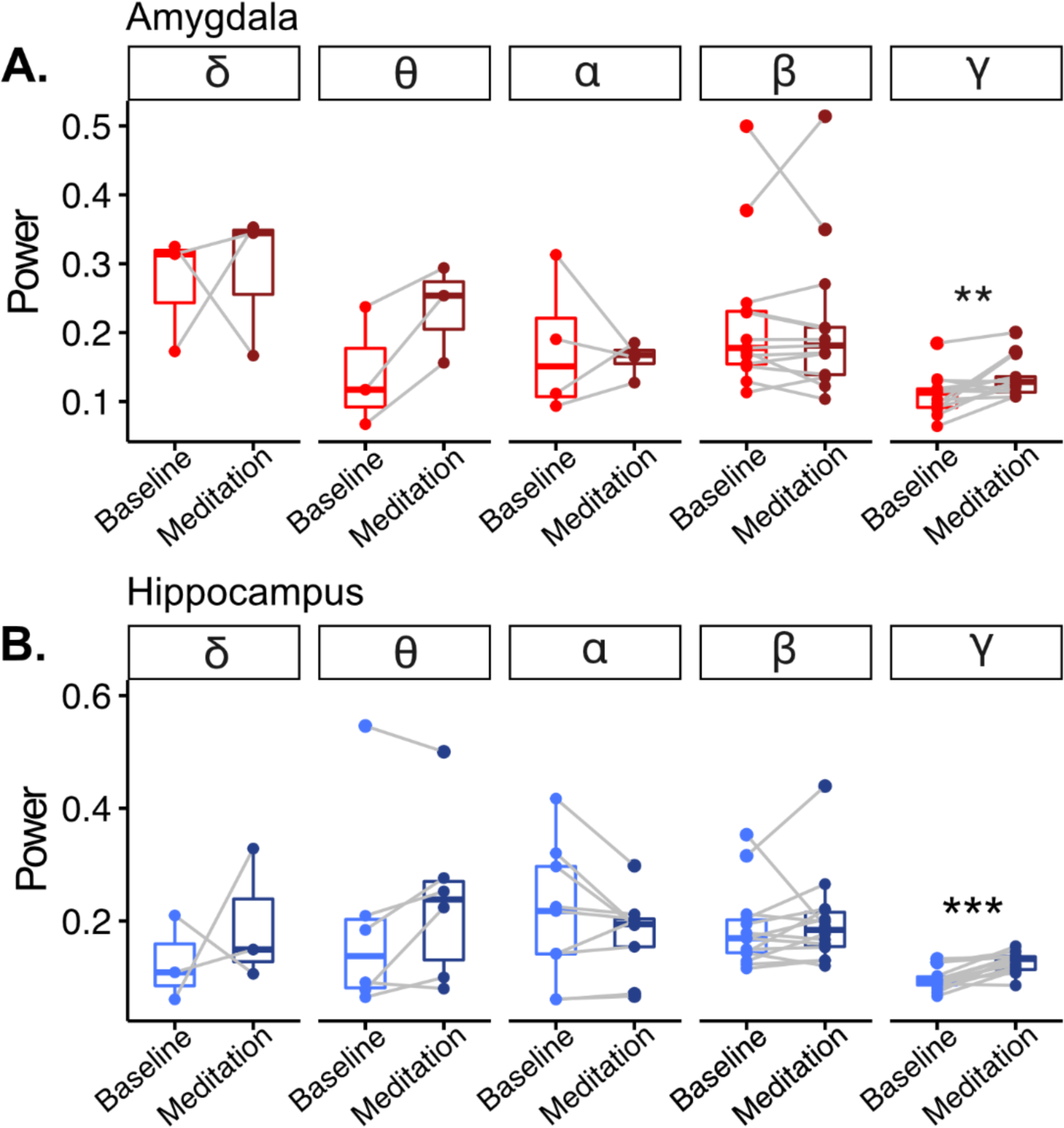
Increased γ oscillatory power in meditation compared to baseline epochs. We examined power modulation across conditions (baseline vs LKM) by averaging spectral power estimates (the periodic component of FOOOF model fit) within frequency bands (δ=2-4, θ=4-8, α=8-13, β=13-30, γ=30-55) and compared across conditions. For this analysis, we employed only the subset of channels that showed significant oscillations in both conditions, which varied across frequency bands. **A. Increased** γ **oscillatory power in amygdala electrodes during meditation.** We observed a significant increase in amygdala γ power (30-55 Hz) during LKM compared to baseline (p<0.01, one-sided paired-samples Wilcoxon signed rank test; all other frequency bands p>0.05). **B. Increased** γ **oscillatory power in hippocampal channels during meditation.** As in the amygdala channels, we observed a significant increase in amygdala γ power (30-55 Hz) during LKM compared to baseline (p<0.001, one-sided paired-samples Wilcoxon signed rank test; all other frequency bands p>0.05).

**Figure 6.**
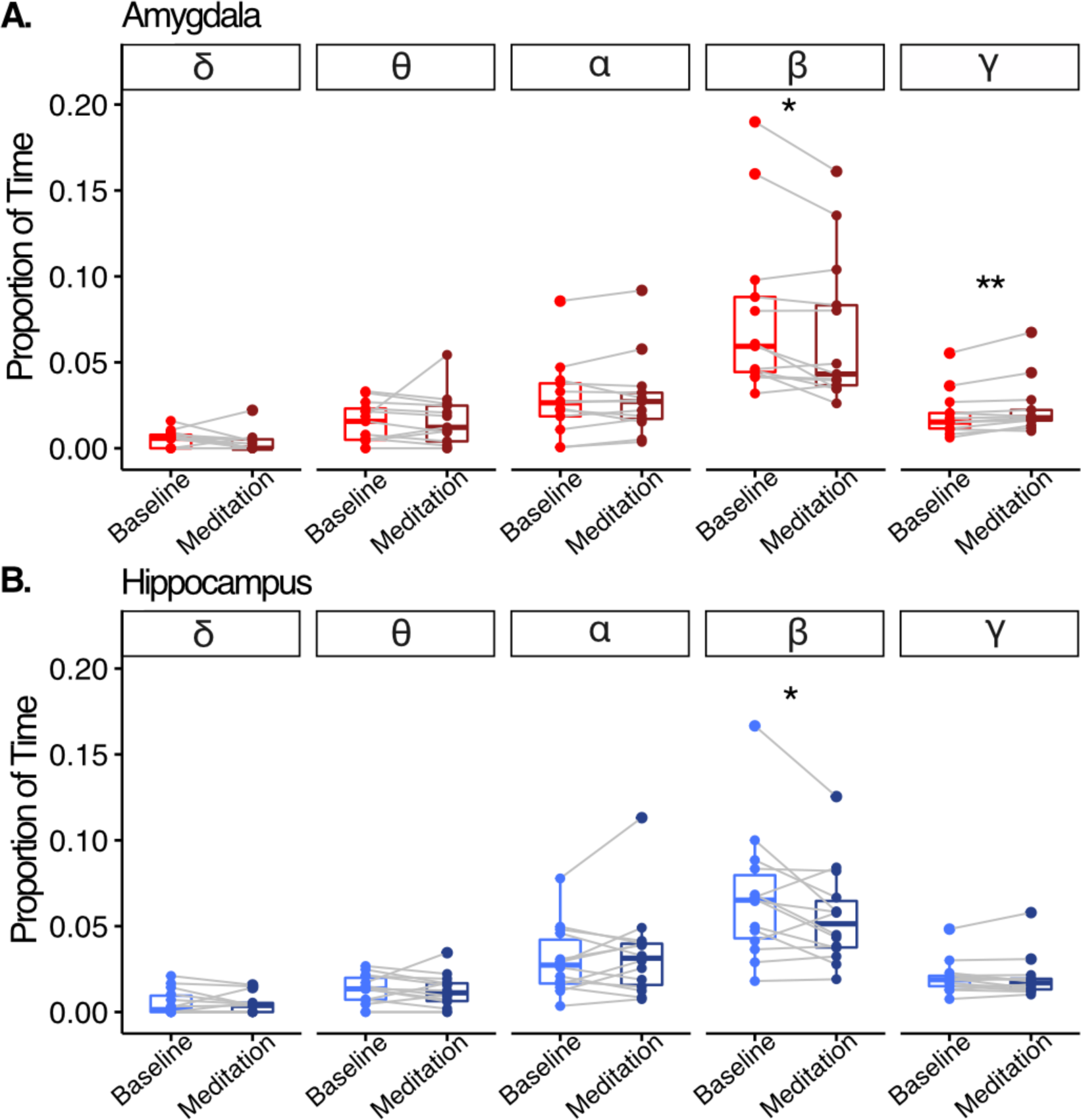
Modulation of oscillatory epochs across meditation states. We used a rhythm detection method (eBOSC) to determine the duration of rhythmic, oscillatory activity. Estimates of oscillatory duration were averaged within frequency bins (δ=2-4, θ=4-8, α=8-13, β=13-30, γ=30-55) and compared between conditions. **A. Amygdala channels showed a decrease in** β **and an increase in** γ **oscillation duration.** The duration of amygdala β-range oscillatory activity during LKM decreased compared to baseline (p < 0.05, one-sided paired-samples Wilcoxon signed rank test), and the duration of amygdala γ-range oscillatory activity during LKM increased compared to baseline (p < 0.01, one-sided paired-samples Wilcoxon signed rank test; all other frequency bands p>0.05). **B. Hippocampus channels showed a decrease in** β **oscillation duration.** The duration of hippocampal β-range oscillatory activity during LKM decreased compared to baseline (p < 0.05, one-sided paired-samples Wilcoxon signed rank test; all other frequency bands p>0.05).

### Amygdala and hippocampus β oscillatory duration decreases during LKM

β oscillations reflect rhythmic phase-locked activity and are thought to facilitate long-range information exchange via cross-regional synchrony. The synchrony afforded by ongoing β oscillations poises neural ensembles to quickly reorganize and integrate incoming information. In human and nonhuman primates, synchrony in lower-frequency (i.e., β) oscillations is thought to modulate attention switches (Li et al., 2017; Miller & Buschman, 2013). This physiological mechanism provides an attentional spotlight that facilitates effective information-gathering from the environment (Buschman & Miller, 2007). This affordance is advantageous in circumstances which require real-time monitoring and adapting to one’s environment. In contrast, one seeks to reduce the influence of external distractions during LKM. Therefore, our present findings of decreased β oscillatory duration during LKM compared to baseline supports the notion that β oscillations are a physiological mechanism for long-range temporal synchrony that regulates attention selection in service of LKM. Transient β bursts, instead of sustained rhythmic oscillations, facilitate cognitive processes through functional inhibition (Lundqvist et al., 2024). β bursts dictate the spatial and temporal basis of activation relevant for memory- and attention-related processing, while gamma power reflects the processing itself ((Brincat et al., 2021). Increased cortical β oscillations have been correlated with negative mood disorders (Kim et al., 2022; Li et al., 2017; Young et al., 2024). Decreased β oscillations with LKM may also represent a shift from negative emotional states, in turn for more positively-salient γ oscillations at the network level.

### Meditation does not affect aperiodic activity or differentially impact the hippocampus versus amygdala

Our finding that meditation was not accompanied by changes in aperiodic neural activity supports that any LKM practice-induced neural fluctuations are specifically related to oscillatory events, not other physiological processes (Fig 3). We did not find differences in any oscillatory metric (periodic/aperiodic FOOOF metrics or eBOSC) between the amygdala and hippocampus. This may indicate they are similarly modulated, with an important caveat that our amygdala contacts are bipolar referenced to the head of the hippocampus. Such confounders can be overcome in future experiments with adjustments of the recording paradigms of the RNS to monopolar, yet, at this time are beyond the scope of this paper.

### Naturalistic study on the neural correlates of meditation

This work on the neural correlates of meditation using RNS has a multitude of advantages over other approaches. Mainly, the RNS allows detailed spatiotemporal study of deep brain structures in a naturalistic setting. While work from fMRI and other neuroimaging studies have gleaned important findings, they are limited in their temporal resolution. Clinically relevant findings from these studies, such as biofeedback for treatment of negative emotions (Zhu et al., 2019), is difficult outside of the naturalistic setting, as with RNS. Our work builds on the uses of RNS to study the neural basis of human brain function in the naturalistic setting, which has been implemented for tasks ranging from spatial navigation (M. Aghajan et al., 2017; Meisenhelter et al., 2019), to speech (Rao et al., 2017).

### Future Directions

These results are the first to document acute intracranial electrophysiology changes with loving kindness meditation. Specifically, we see an increase in γ oscillations and a decrease in β burst duration in the amygdala and hippocampus with LKM. While we observe acute changes in amygdala and hippocampal electrophysiology during LKM, we did not assess long-term changes that may occur with LKM practice. The meditation literature has defined “state changes” as temporary modulations of mind (and implicitly brain function) that occur with meditation, versus trait changes as long-term, enduring effects of meditation on the mind (and thus brain) (May et al., 2011). Our research documents brain signals that may contribute to the acute state changes of LKM but does not explore long-term brain changes that may underlie traits unique to long-term meditators. Yet, the RNS system enables long-term tracking of electrophysiology, which may be applied over the course of a novice meditator’s practice as they journey to a higher level of expertise. Further, alternative forms of meditation, such as mindfulness, may be similarly assessed to determine the varying effects of meditation practice on brain state and trait changes. The work presented here focuses on hippocampal and amygdala electrophysiology, as these are the typical targets of RNS implantation for epilepsy. Complementary studies on intracranial EEG recordings in patients implanted with a wider range of contact locations for seizure localization (sEEG) may provide additional insight into the circuit-level activity changes associated with meditation. We feel both approaches are complementary and will improve future investigations via a feedback loop. There thus remains an exciting opportunity for exploration on how meditation practice exerts acute state changes and long-term trait changes through modulation of intracranial electrophysiology in the human brain.

### Summary

Our findings reveal increased hippocampal and amygdala γ power associated with LKM, and amygdala-specific increases in γ oscillation duration. Further, we identified a global decrease in amygdala ad hippocampus β oscillatory duration during LKM. These findings provide novel, anatomically localized and neurophysiologically detailed for the role of these regions in meditation, and suggest an association with their known roles in memory and emotional regulation processes. In addition, they confirm the potential of first-time LKM practice to induce transient physiological alterations in brain regions whose maladaptive functioning is implicated in mental health disorders. By identifying physiological mechanisms that are amenable to noninvasive neuromodulation via LKM, we highlight LKM’s potential as a targeted therapeutic intervention.

## MATERIALS AND METHODS

### RNS participants

Eight DRE patients participated in the present study (n= 3 female, mean age = 44.75±7.67 years). All participants had been previously implanted with the NeuroPace RNS System to treat DRE. Electrode placement was solely determined by clinical criterion. A structured clinical interview was conducted based on the NIH Common Data Elements Battery for Epilepsy Patients prior to participation. Participants provided informed consent for participating in the present study which was approved by the Mount Sinai Institutional Review Board (IRB).

### RNS data acquisition

RNS is an FDA-approved chronically implanted invasive intervention to treat patients with DRE by continuously monitoring for abnormal electrical activity and delivering electrical stimulation to circumvent seizure onset (Fig 1A). RNS devices provide access to intracranial LFP activity from the stimulating electrode in post-surgical, chronic conditions, which provides an opportunity to record intracranial LFP data from deep brain regions. iEEG storage in the RNS Neurostimulator was manually triggered by the experimenter to mark the start and end of the experimental session. During the experimental session, the RNS Neurostimulator continuously recorded iEEG activity in two (unilateral implantation) or four (bilateral implantation) bipolar channels depending on each patient’s electrode placement. Bipolar derivations were carried out between the two most anterior and the two most posterior channels, typically implanted along the anterior-posterior axis of the amygdala-hippocampus (Fig 2A). iEEG activity was continuously recorded from all available bipolar channels across two depth electrode leads in each patient at a sampling frequency of 250 Hz.

### Behavioral task

The experimental session consisted of a 15-minute audio-guided meditation paradigm. All experimental sessions occurred in Mount Sinai’s Quantitative Biometrics Laboratory (Q-Lab) which was designed to emulate Japanese teahouses and gardens, affording patients a restorative environment for participation (Fig 1B). The present study’s LKM paradigm was an audio-only pre-recording to ensure standardization across participants. The recording utilized material from the Healthy Minds Program and Mindfulness-Based Stress Reduction course through Palouse Mindfulness. Both programs provide open-source, empirically-evidenced LKM resources (Dahl et al., 2015). The recording was performed by a neuropsychologist with specialist training Acceptance and Commitment Therapy, a psychological intervention that utilizes mindfulness, and experience in facilitating meditation sessions for epilepsy patients. An experimenter was present for each experimental session to monitor potential seizure activity. The meditation paradigm began with 5-minutes of audio-guided instruction during which participants passively listened to discussion of meditation’s objective and were guided in best practices for engaging effectively in LKM. This period was considered baseline in further analysis. Instruction was followed by 10-minutes of LKM during which participants were guided to wish well to others by repeating phrases silently, such as “may you be happy, may you be free of pain”. The meditation builds from wishing happiness to loved ones before moving on to acquaintances, and then to those the practitioner feels neutral towards or even actively dislikes. Through active attention regulation, practitioners to exert top-down attention on a target object while disengaging attentional resources from irrelevant objects. The goal-directed modulation of attention and its neural underpinnings has been well-characterized in humans (Leong et al., 2017). Until now, significant technical limitations have prevented the direct recording of neural activity with the spatial and temporal resolution necessary to potentially detect neurophysiological fluctuations induced by meditative states, especially in novice meditators. RNS patients present a unique opportunity to record intracranial electrophysiology (iEEG) in freely behaving, post-operative humans. For these reasons, we chose to focus the present iEEG study on LKM. Participants remained seated for the duration of the experimental session. For analysis purposes, the first two minutes of instruction were selected as the baseline epoch, and two minutes in the middle of meditation during each patient’s experimental session were selected as the meditation epoch for comparison. These conditions were referred to as baseline and meditation, respectively.

### Analysis

Analyses were performed with custom MATLAB (electrode localization), Python (iEEG preprocessing and quantification), and R (statistics and visualization).

### Electrode localization and iEEG preprocessing

Electrode localization was performed with MATLAB LeGUI. We co-registered each participant’s post-operative CT image to their pre-operative whole brain MRI. All electrodes determined to be in the amygdala or hippocampus following localization were included in further analysis (amygdala: electrode, bipolar channels; hippocampus: electrodes, bipolar channels). The anatomical location of each electrode was determined using the Yale Brain Atlas (McGrath et al., 2022), a whole-brain atlas of the human cortex, hippocampus, amygdala created using 3866 electrodes across 25 iEEG subjects, and verified through visual examination. Epileptic or noisy activity was manually removed following visual inspection of each channel similar to previously published methods (Saez et al., 2018). Similar to previous results (M. Aghajan et al., 2017; Maoz et al., 2023; Saez et al., 2018), we excluded ∼6% of the data from further analyses due to the presence of epileptic or noisy activity. The proportion of excluded data across amygdala and hippocampus channels was not significantly different between conditions (p > 0.05 for all channels, Pearson’s chi-square goodness-of-fit test).

### Quantifying oscillatory prevalence, power, and duration

We computed the power spectral density (PSD) of the iEEG signal between 0 and 125 Hz using the Welch method (2s-segments, 12.4% overlap).

### Fitting Oscillations and 1/f (FOOOF) Method

PSDs for each channel across conditions were analyzed using the FOOOF method (Donoghue et al., 2020) which parameterizes aperiodic and periodic features of the power spectrum allowing us to determine whether FA meditation-induces true oscillatory changes in neural activity or impacts other neurophysiological processes. Consistent with prior work leveraging the FOOOF method to assess human electrophysiological data, we visually inspected each PSD to determine the appropriate method of aperiodic fitting (linear versus nonlinear). Across most channels (n=26), PSDs depicted a bend in higher frequencies (>40 Hz), therefore spectral parameterization with FOOOF was performing using a knee to appropriately capture these power spectrums’ nonlinear dynamics (Fig 2B). This was consistent with existing human electrophysiology literature, especially when evaluating higher frequency domains (> 40 Hz). In one channel, a bend was not observed in the PSD. Spectral parameterization for this channel was performed without a knee. Using fitted exponent and knee parameters for each channel, we calculated the knee frequency to evaluate group differences. The knee frequency reflects the frequency in which the aperiodic slope of the PSD changes. Further, peak width was restricted to 1-8 Hz to minimize overfitting. Model fit was evaluated according to R-squared and error metrics (Amygdala: median R2(median error) = baseline (0.99(0.03)), meditation (0.99(0.04)); Hippocampus: baseline (0.99(0.03)), meditation (0.99(0.03))). R-squared for both regions was in accordance with existing literature (Kopčanová et al., 2024) and not significantly different between conditions suggesting good fit for both the amygdala (p > 0.05, two-sided paired-samples Wilcoxon signed rank test) and hippocampus (p > 0.05, two-sided paired-samples Wilcoxon signed rank test).

Using the FOOOF method, we fit the aperiodic (offset, knee, exponent) and periodic (power, bandwidth, center frequency) components of each iEEG channel’s power spectrum between 2 and 55 Hz for each condition (Fig 2B). This allowed us to separate true oscillatory features of the PSD versus background (1/f) neural activity, characterized by the aperiodic components. The aperiodic offset, or broadband intercept, reflects the up/down translation of the spectrum while the aperiodic exponent reflects the overall curve of the aperiodic component with a smaller exponent reflecting a shallower power spectrum (Fig 2B). Oscillations are reflected as narrowband peaks in power above the aperiodic component of the PSD. Parameterizing both the periodic and aperiodic components of the PSD allows us to compare differences in the presence/absence of oscillations at individual frequency bands between regions and conditions, as well as the periodic features of those oscillations while accounting for the background 1/f aperiodic profile which reflects other physiological processes. Periodic components of the power spectra refer to activity with a characteristic frequency. Of interest for the present study is frequency-specific power, which quantifies the amount of energy contained in the iEEG signal at a particular frequency band. Aperiodic components of the power spectra refer to recorded activity with no characteristic frequency. We assessed LKM-associated differences in aperiodic offset, which measures the overall up/down translation of the power spectra, exponent, which reflects the slope of the aperiodic component of the power spectra, and the knee frequency, which quantifies the aperiodic component’s “bend” or transition point wherein the dominant characteristics of the neural signal change (Fig 2B; Donoghue et al., 2020). For each channel, we estimated power in putative frequency bands by taking the average power of peaks above the aperiodic component falling within a priori designated ranges according to their center frequency (δ=2-4Hz, θ=4-8Hz, α=8-13Hz, β=13-30Hz, γ=30-55Hz).

### Extended Better Oscillation Detection (eBOSC) Method

To assess FA meditation-induced temporal differences in oscillatory activity, we estimated the duration of oscillations by applying the eBOSC method (Hu et al., 2014; Kosciessa et al., 2020). By calculating both the strength (i.e., amplitude) and duration of a given oscillation, eBOSC provides a comprehensive characterization of oscillatory events. eBOSC allowed us to estimate the duration of oscillations by identifying significant oscillation that surpass a statistical power threshold, extracting the number of detected cycles in each significant oscillation, and converting the number of detected cycles to duration in a frequency-specific manner. eBOSC first performs time-frequency decomposition on continuous data. The resulting power spectra are fit linearly using robust regression to estimate the relationship between log(frequency) and log(power). A power threshold for each frequency was set at the 95% of a χ^2^-distribution of power values centered around the fitted estimate of background power at a given frequency based on the linear model fit. This establishes a statistical power threshold for determining a significant oscillation at a given frequency. Using the derived power threshold, oscillations and their associated time intervals were extracted. The duration threshold was set to zero, allowing us to compare both transient and sustained oscillatory episodes. The proportion of time spent oscillating at each frequency is calculated by summing detected time intervals and normalizing by total duration of a given iEEG recording.

### Statistics

#### Data quality checks

We used a Nonparametric Pearson’s chi-square goodness-of-fit test to determine whether there was a significant difference in the proportion of data within each condition following pre-processing. We used a Nonparametric Wilcoxon rank-sum test to check whether FOOOF model fit (R^2^) differed significantly between conditions.

#### Hypothesis testing

We used a Nonparametric Fisher’s exact test to assess whether a nonrandom association exists between experimental condition (baseline versus meditation) and categorical variables (presence versus absence of an oscillation within given frequency band). To assess significant differences between conditions for linear variables (oscillatory power and duration), we used Nonparametric Wilcoxon rank-sum tests. One-sided Wilcoxon rank-sum test were implemented where applicable based on empirical literature regarding meditation-induced modulation of β and γ oscillations (Irrmischer, Houtman, et al., 2018; Irrmischer, Poil, et al., 2018; Lutz et al., 2004)

#### Data and Software Availability

Anonymized data from participants who consented to data sharing is available upon request with the approval of NeuroPace. Analysis scripts can be found at the following GitHub repository: https://github.com/christinamaher/iEEG_LKM

## Acknowledgements

We thank Jill Gregory, MFA, CMI, Associate Director of Instructional Technology at the Icahn School of Medicine at Mount Sinai, for illustrating Figure 1A. We would like to thank the patient and research and surgical staff at the recording site for their support and dedication.

## SUPPLEMENTARY FIGURES

**Supplementary Figure 1.**
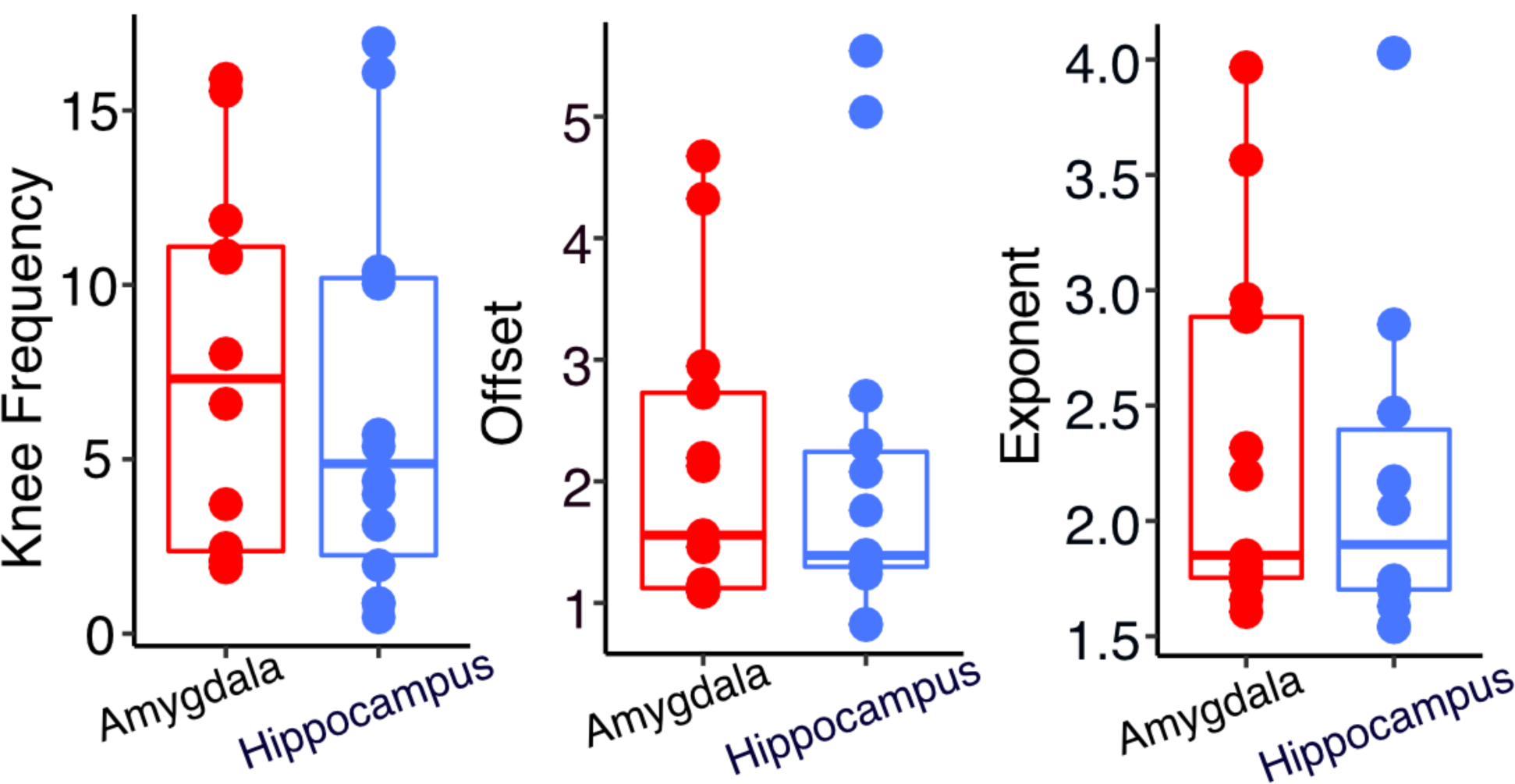
No difference in aperiodic neural components between amygdala and hippocampus. The aperiodic component of FOOOF model fit (knee frequency, offset, and exponent) from baseline epochs were extracted and compared between regions (amygdala and hippocampus). No significant differences in knee frequency, offset, or exponent of FOOOF model fit between amygdala electrodes (n=13 bipolar channels) and hippocampus (n= 14 bipolar channels) at baseline (all p>0.05, Two-sided sample Wilcoxon signed-rank test).

**Supplementary Figure 2.**
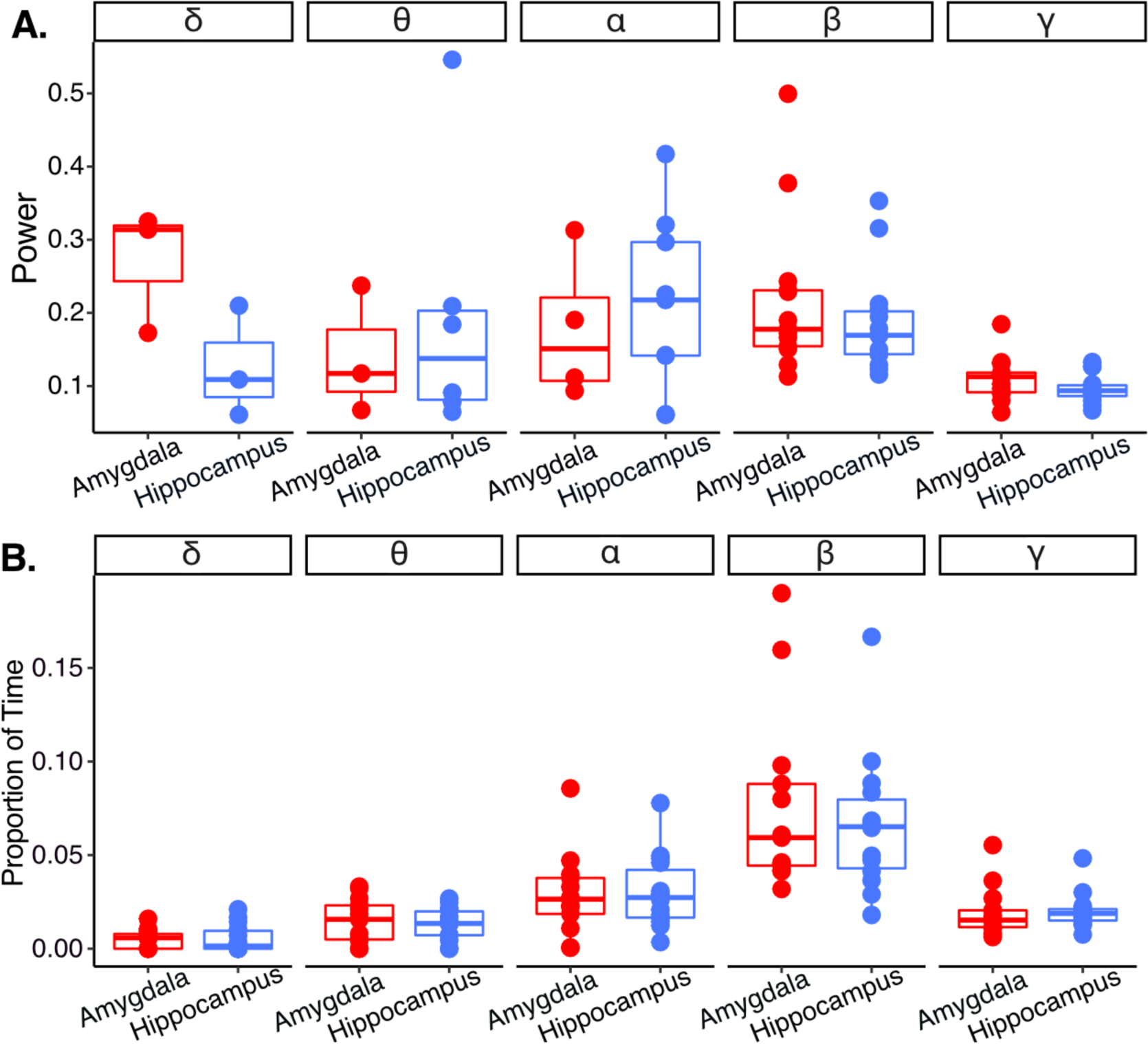
Comparison of baseline oscillatory power and duration between amygdala and hippocampus. We examined differences in oscillatory power and duration between amygdala and hippocampus channels at baseline. **A. No difference in baseline oscillatory power between amygdala and hippocampus.** We examined power modulation during baseline epoch between regions by averaging spectral power estimates (the periodic component of FOOOF model fit) within frequency bands (δ=2-4, θ=4-8, α=8-13, β=13-30, γ=30-55) and compared between regions. For this analysis, we employed only the subset of channels that showed significant oscillations in both experimental conditions, which varied across frequency bands. We did not observe significant differences in power across frequency bands between regions (all p>0.05, Two-sided sample Wilcoxon signed-rank test). **B. No difference in baseline oscillatory duration between amygdala and hippocampus.** We used a rhythm detection method (eBOSC) to determine the duration of rhythmic, oscillatory activity during baseline epoch. Estimates of oscillatory duration were averaged within frequency bins (δ=2-4, θ=4-8, α=8-13, β=13-30, γ=30-55) and compared between regions. We did not observe significant differences in duration across frequency bands between regions (all p>0.05, Two-sided sample Wilcoxon signed-rank test).

**Supplementary Figure 3.**
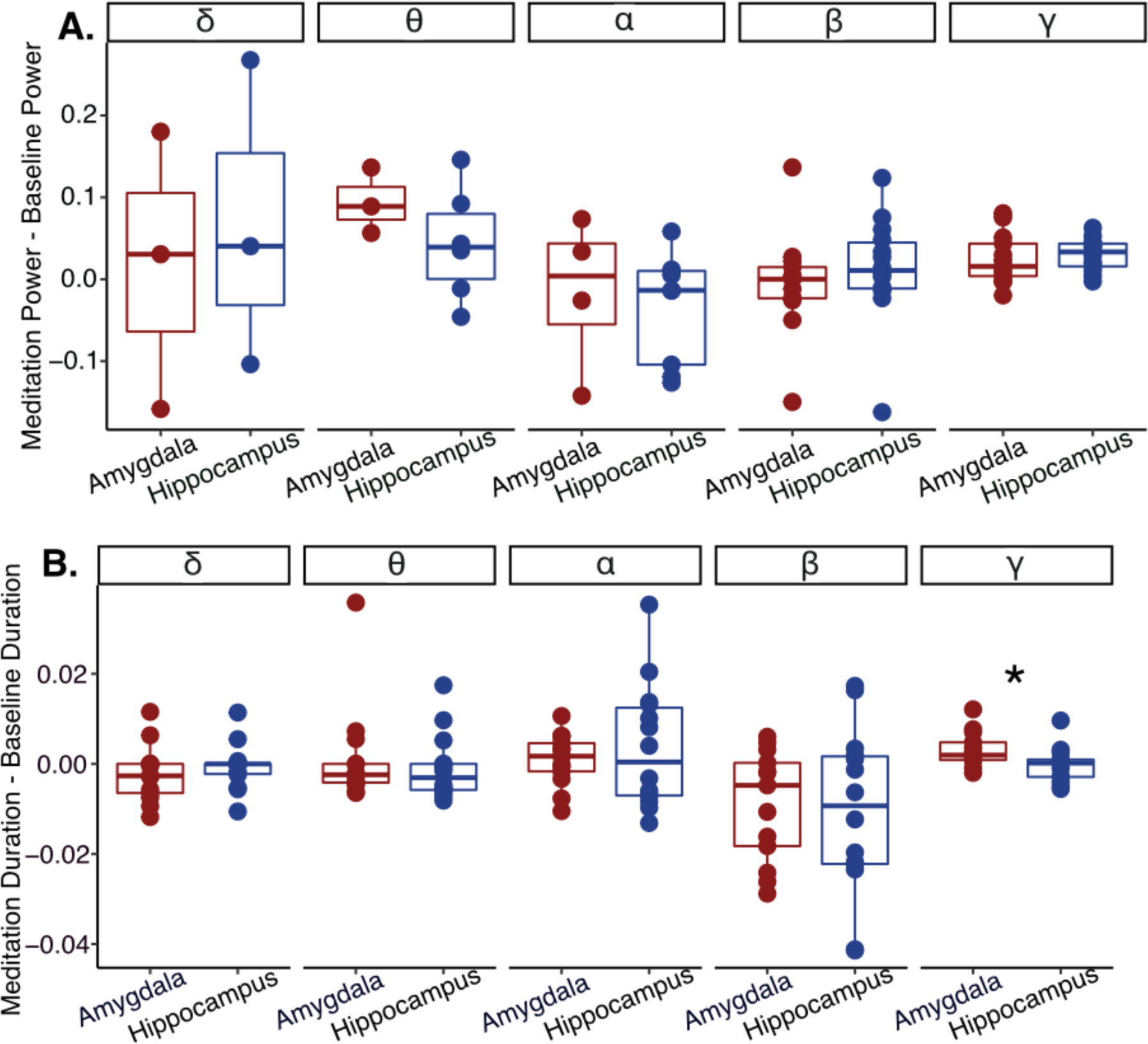
Comparison of condition-dependent change in oscillatory power and duration between amygdala and hippocampus. We examined differences in change in oscillatory power and duration between amygdala and hippocampus channels between conditions. **A. No difference in oscillatory power changes between baseline and meditation between amygdala and hippocampus.** We examined change in oscillatory power by subtracting power at different frequency bands at baseline from power observed during meditation (meditation epoch power – baseline epoch power). We did not observe significant differences in the change in power across frequency bands between regions (all p > 0.05, Two-sided sample Wilcoxon signed-rank test). **B. Amygdala channels showed a larger increase in γ oscillation duration from baseline to meditation compared to hippocampus channels.** We examined change in oscillatory duration by subtracting proportion of time oscillatory bursts were detected at different frequency bands during the baseline epoch from the duration of oscillatory bursts observed during the meditation epoch (meditation epoch duration – baseline epoch duration). The change in duration of γ oscillations from baseline epoch to meditation epoch was significantly greater in the amygdala compared to hippocampus (p < 0.05, Two-sided sample Wilcoxon signed-rank test; all other frequency bands p>0.05).

